# Bile acid metabolism is altered in multiple sclerosis and supplementation ameliorates neuroinflammation

**DOI:** 10.1101/627356

**Authors:** Pavan Bhargava, Leah Mische, Matthew D. Smith, Emily Harrington, Kathryn C Fitzgerald, Kyle Martin, Sol Kim, Arthur Anthony Reyes, Jaime Gonzalez-Cardona, Christina Volsko, Sonal Singh, Kesava Varanasi, Elias S. Sotirchos, Bardia Nourbakhsh, Ranjan Dutta, Ellen M. Mowry, Emmanuelle Waubant, Peter A. Calabresi

## Abstract

Multiple sclerosis (MS) is an inflammatory demyelinating disorder of the CNS. Bile acids are cholesterol metabolites that can signal through receptors on cells throughout the body, including the CNS and immune system. Whether bile acid metabolism is abnormal in MS is unknown. Using global and targeted metabolomic profiling, we identified lower levels of circulating bile acid metabolites in multiple cohorts of adult and pediatric MS patients compared to controls. In white matter lesions from MS brain tissue, we noted the presence of bile acid receptors on immune and glial cells. To mechanistically examine the implications of lower levels of bile acids in MS, we studied the *in vitro* effects of an endogenous bile acid – tauroursodeoxycholic acid (TUDCA) on astrocyte and microglial polarization. TUDCA prevented neurotoxic (A1) polarization of astrocytes and pro-inflammatory polarization of microglia in a dose-dependent manner. TUDCA supplementation in experimental autoimmune encephalomyelitis reduced severity of disease, based on behavioral and pathological measures. We demonstrate that bile acid metabolism is altered in MS; bile acid supplementation prevents polarization of astrocytes and microglia to neurotoxic phenotypes and ameliorates neuropathology in an animal model of MS. These findings identify dysregulated bile acid metabolism as a potential therapeutic target in MS.

## Introduction

Bile acids are end-products of cholesterol metabolism that have multiple biological functions besides aiding lipid absorption in the gut^1^. Primary bile acids are mainly produced in the liver, but other tissues are capable of bile acid synthesis through an alternative “acidic” pathway. Primary bile acids are modified (conjugated with glycine or taurine) and then stored in the gall bladder until they are secreted into the gut, where they are further modified through enzymes present in gut bacteria to produce secondary bile acids (Figure 1a) ^1^. The majority of bile acids are reabsorbed and undergo enterohepatic recirculation through the portal venous system with a small fraction reaching the systemic circulation. Circulating bile acids can act on receptors, both nuclear (Farnesoid X receptor - FXR) and cell surface (G-protein coupled bile acid receptor1 - GPBAR1), found on cells including those in the brain and the immune system^2,3^. While the role of altered bile acid metabolism has been extensively explored in metabolic diseases, only recently were abnormalities described in neurological disease^3,4^. In a large cohort of Alzheimer’s disease (AD) patients, abnormal circulating bile acid metabolite levels predicted worse outcomes and were hypothesized to be related to alteration of the gut microbiota^4^.

**Fig. 1.**
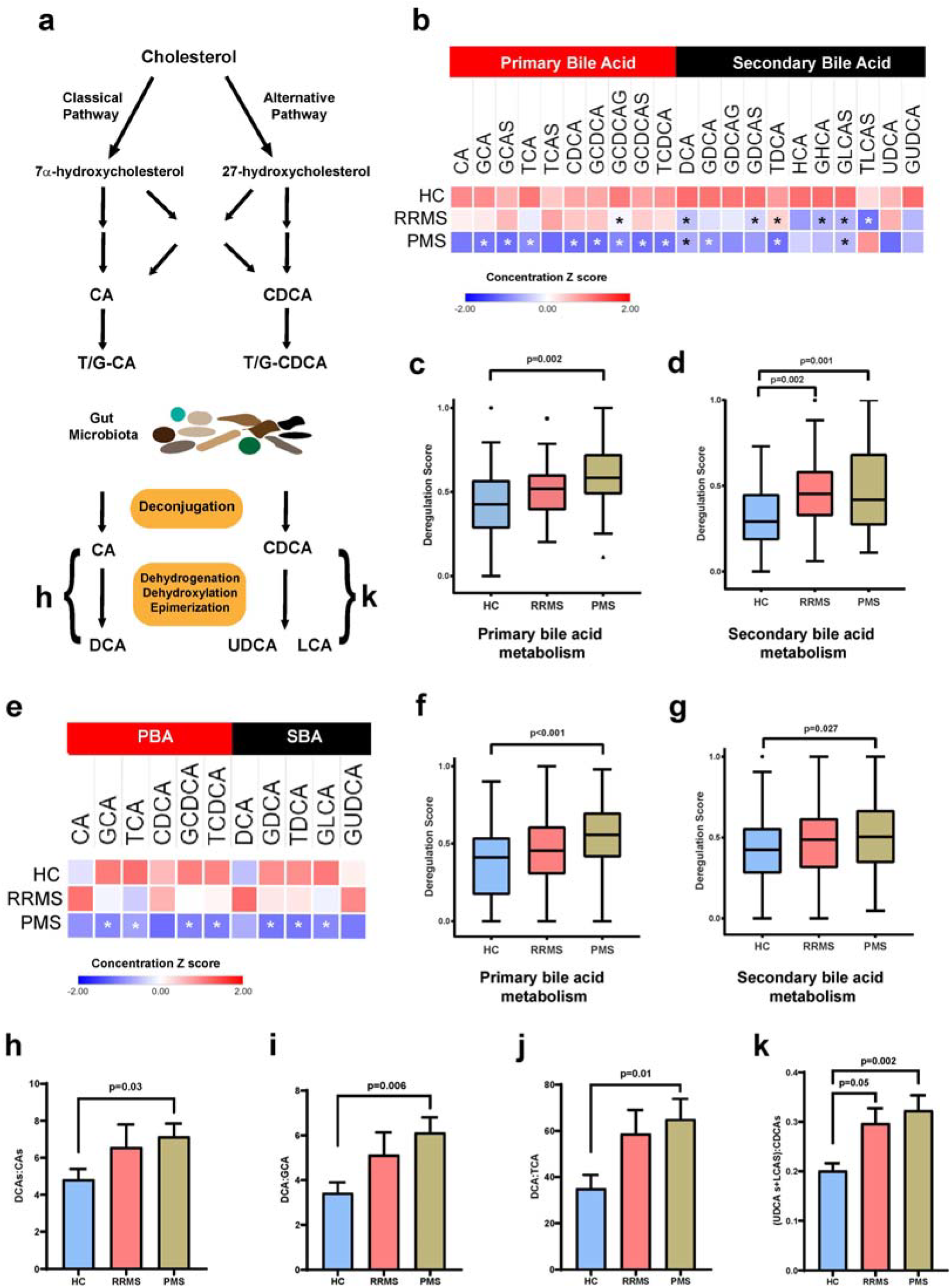
Bile acid metabolism is altered in Multiple Sclerosis. (**a**) Overview of bile acid metabolism. CA – cholic acid, CDCA – chenodeoxycholic acid, G/T – glycine/ taurine conjugated, DCA – deoxycholic acid, LCA – lithocholic acid, UDCA – ursodeoxycholic acid. (**b**) Heat map of mean standardized bile acid metabolite concentrations derived from untargeted metabolomic profiling in the discovery cohort – RRMS (n=56), PMS (n=52) and healthy controls (n=50). Multiple bile acid metabolites in both primary and secondary bile acid metabolism pathways are lower in the MS groups compared to controls. Asterisks denote significant differences compared to healthy controls based on multivariate linear regression models adjusting for age, sex and race (p<0.05). **(c)** Box-plots of pathway deregulation scores for primary bile acid metabolism i the discovery cohort demonstrate significant abnormality in the PMS group compared to controls. For all box plots -center line – median, box – 25^th^ and 75^th^ percentiles, whiskers – 1.5 × interquartile range and points – outliers. **(d)** Box-plots of pathway deregulation scores for secondary bile acid metabolism in the discovery cohort demonstrate significant abnormality in the RRMS and PMS groups compared to controls (p-values for c and d derived from multivariate linear regression models adjusting for age, sex and race). **(e)** Heat map of mean standardized bile acid metabolite concentrations derived from targeted metabolomic profiling in the validation cohort – RRMS (n=50), PMS (n=125) and healthy controls (n=75). Multiple bile acid metabolites in both primary and secondary bile acid metabolism pathways are lower in the MS groups compared to controls. Asterisks denote significant differences as in panel b. (**f)** Box-plot of pathway deregulation scores for primary bile acid metabolism in the validation cohort demonstrate significant abnormality in the PMS group compared to controls. **(g)** Box-plots of pathway deregulation scores for secondary bile acid metabolism in the validation cohort demonstrate significant abnormality in the PMS group compared to controls (p-value is derived as in c and d). **(h)** Increased DCA (sum of all DCA metabolites) to CA (sum of all CA metabolites) metabolite ratio was noted in the PMS group compared to controls. The ratios of DCA to individual conjugated CA metabolites were also higher in the PMS group **(i and j)**. **(k)** The ratio of the sum of UDCA and LCA metabolites to the sum of CDCA metabolites was higher in the PMS group compared to controls. Bars in h-k represent mean with error bars representing s.e.m and p-values are derived from a linear regression model.

Multiple sclerosis (MS) is a chronic inflammatory demyelinating disorder and both genetic and environmental factors play a role in its etiopathogenesis^5–7^. Gut dysbiosis has been described in MS patients with altered abundance of several bacterial species involved in bile acid metabolism^8–10^. In experimental autoimmune encephalomyelitis (EAE), neuroinflammation is associated with altered cholesterol metabolism in astrocytes^11^ and abnormalities in circulating bile acid metabolites^12^ providing further rationale for their evaluation in MS. An FXR agonist led to amelioration of disease in EAE, through anti-inflammatory effects on myeloid cells^13,14^, while GPBAR1 activation produced anti-inflammatory effects in innate immune cells and glial cells^15,16^. Since bile acids have important effects on key inflammatory pathways and can be neuroprotective^17,18^, we hypothesized that bile acid metabolism may play a role in modulating neuroinflammation in MS.

## Results

### Bile acid metabolism is altered in adult-onset multiple sclerosis

We performed global metabolomic profiling on plasma from a discovery cohort consisting of adult patients with relapsing-remitting MS (RRMS, n=56), progressive MS (PMS, n=51) or healthy controls (n=52) and identified 25 bile acid metabolites (Supplementary Table 1). As expected, RRMS patients were younger and PMS patients were older than the healthy controls (Supplementary Table 2). Lower levels of multiple primary bile acid metabolites were seen in the PMS group compared to controls (Figure 1b. Supplementary Figure 1a). Comparison of pathway deregulation scores between groups adjusting for age, sex and race revealed significantly higher scores (denoting greater abnormality) for primary bile acid metabolism in the PMS group (Figure 1c). Lower levels of multiple secondary bile acid metabolites were seen in both RRMS and PMS compared to healthy controls (Figure 1b, Supplementary Figure 1b). Secondary bile acid metabolism pathway deregulation scores (adjusted for age, sex, and race) were higher for both MS groups compared to controls (Figure 1d).

To extend our initial observations, we performed targeted metabolomic profiling of 15 bile acids (Supplementary Table 3) in plasma from a larger cohort of MS patients (RRMS: n=50, PMS: n=125) and controls (n=75). Again, the PMS patients were older than RRMS patients and healthy controls (Supplementary Table 2). Lower levels of multiple primary and secondary bile acids were seen in the PMS group compared to controls (Figure 1e, Supplementary Figure 1c and d). Comparison of between-group pathway deregulation (adjusting for age, sex, and race) revealed significantly higher scores (denoting greater abnormality) for both primary and secondary bile acid metabolism in the PMS group compared to controls (Figure 1f and g).

To model metabolism of bile acids in the gut, we studied the ratio of secondary bile acid metabolites to their primary bile acid precursors, since secondary bile acid production requires multiple reactions catalyzed by enzymes present in the gut microbiota^4,19^. We noted a significant increase in the ratio of deoxycholic acid (DCA) metabolites (DCA, GDCA, TDCA) to cholic acid (CA) metabolites (CA, GCA, TCA) in the PMS group with a trend towards a higher ratio in the RRMS group (Figure 1h). Individual ratios of DCA to either glycine or taurine-conjugated forms of CA were also elevated in the PMS (Figure 1i and j) but not the RRMS group. In the alternative bile acid synthesis pathway, a similar change was seen with an increased ratio of the lithocholic acid (LCA) and ursodeoxycholic acid (UDCA) metabolites to chenodeoxycholic acid (CDCA) metabolites in the PMS but not the RRMS group compared to controls (Figure 1k). Overall, the data from both metabolomics approaches in the adult cohorts revealed alterations in bile acid metabolism in patients with MS, with greater abnormality noted in the PMS group.

### Bile acid metabolism is altered in pediatric-onset multiple sclerosis

We then performed global metabolomic profiling in a cohort of pediatric-onset MS patients (n=31) and age-, race- and sex-matched controls (n=31) drawn from the UCSF Pediatric MS Center (Supplementary Table 4). Lower levels of several primary and secondary bile acid metabolites were seen in the pediatric MS patients compared to controls (Fig 2a-c, Supplementary Figure 2). Several of the metabolites that were differentially altered between groups overlapped with the adult cohorts. Comparison of pathway deregulation scores for primary bile acid metabolism revealed greater abnormality in the pediatric MS group compared to controls (Figure 2d). A significant increase in the ratio of DCA to CA metabolites was found, similar to that seen in adult MS patients (Figure 2f). These results demonstrate that in a cohort of pediatric-onset MS patients derived from an independent site, we identified bile acid metabolism abnormalities similar to those noted in adult MS patients.

**Fig. 2.**
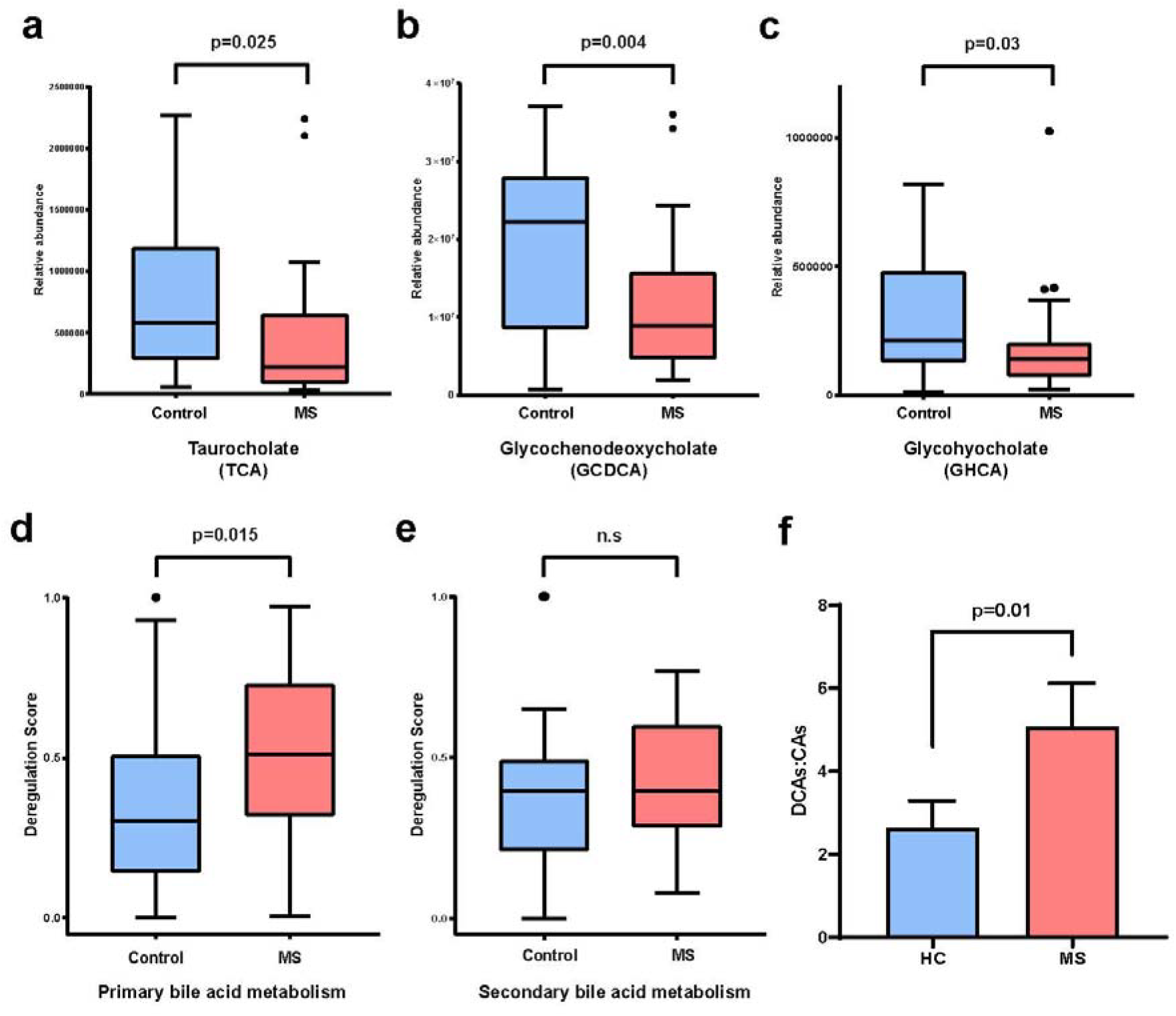
Bile acid metabolism is altered in pediatric-onset Multiple Sclerosis. **(a-c)** Box plots of individual bile acid metabolites in pediatric-onset MS and control groups (n=31 each) demonstrating lower metabolite levels in the MS group. **(d)** Box plots of pathway deregulation scores for primary bile acid metabolism in pediatric-onset MS patients and controls, demonstrating significant abnormality in the metabolic pathway in pediatric MS. **(e)** Box plots of pathway deregulation scores for secondary bile acid metabolism in pediatric-onset MS patients and controls. **(f)** Increased ratio of DCA (sum of all DCA metabolites) to CA (sum of all CA metabolites) metabolites was noted in the MS group. For all box plots center line – median, box – 25^th^ and 75^th^ percentiles, whiskers – 1.5 × interquartile range and points – outliers. The p values for a-are derived from multivariate linear regression models adjusting for age, sex, and race. In f error bars represent standard error of the mean and p value is derived from a two-tailed unpaired Student’s t-test.

### Bile acid receptors are expressed in white matter lesions from MS patients

After establishing altered bile acid metabolism in both adult and pediatric MS patients, we enquired whether the key receptors associated with bile acid signaling are present in MS brain lesions. While a previous study demonstrated the presence of the farnesoid X receptor (FXR) in inflammatory cells in active lesions from RRMS patients^13^, the expression of bile acid receptors has not been studied in lesions from PMS patients. In white matter lesions from PMS autopsy tissue (Figure 3a), FXR positive cells were present in the center (Figure 3a – red box, Figure 3b) of and at the edge (Figure 3a, blue box, Figure 3c) of lesions. The expression of FXR was similar to the distribution and nuclear localization within macrophages (CD68+ cells) as reported previously^13^. Staining within white matter lesions also revealed the presence of GPBAR-1, a cell-surface bile acid receptor, on cells with morphology suggestive of astrocytes (Figure 3d and e) as well as vascular structures (Figure 3d and f). Double immunostaining confirmed the presence of GPBAR-1 staining in cells positive for GFAP confirming that astrocytes in MS lesions express cell-surface bile acid receptors (Figure 3g-i).

**Fig. 3.**
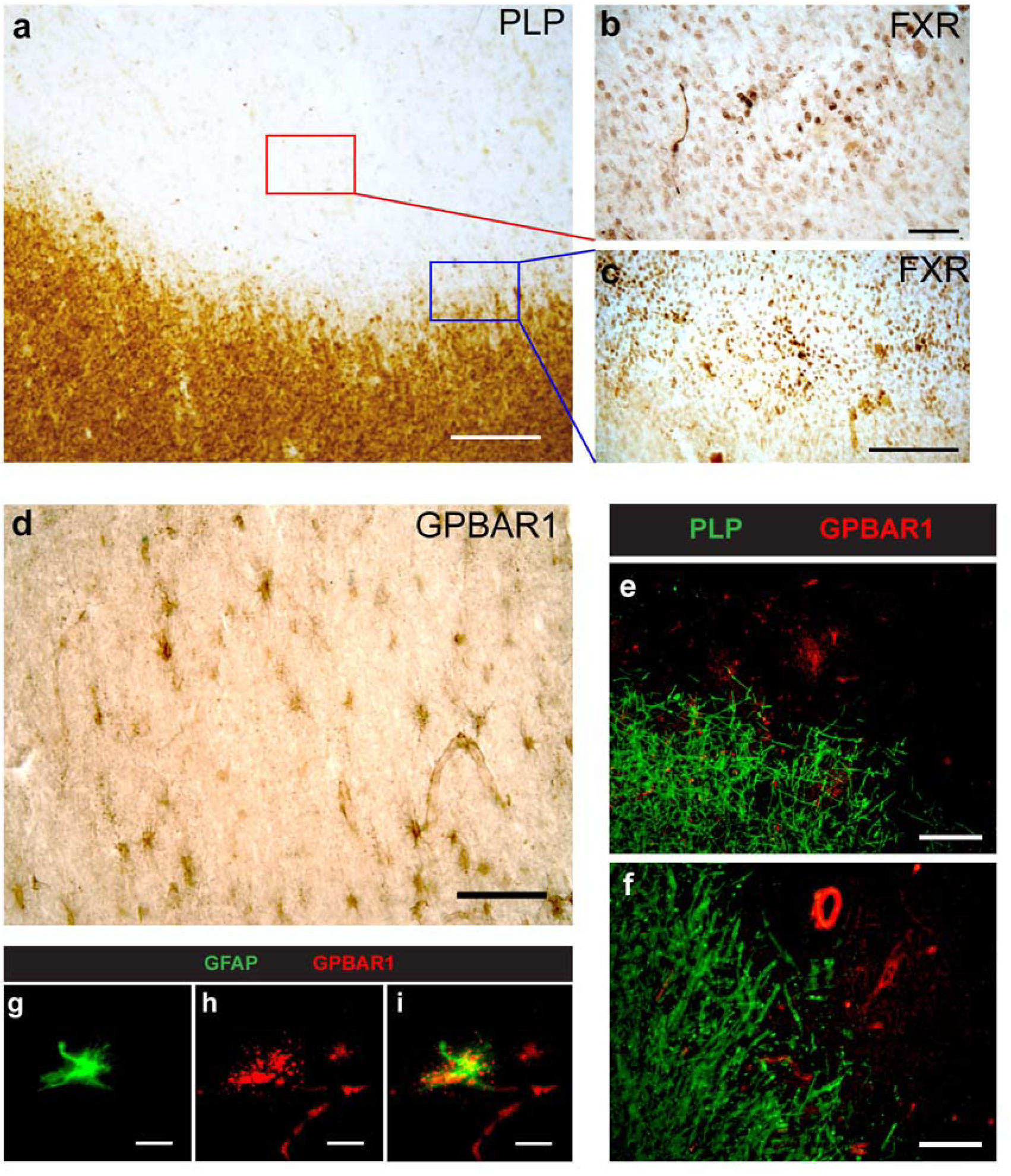
Bile acid receptors are detected in demyelinating lesions in MS brains. (**a**) Immunohistochemistry for proteolipid protein (PLP) identifies a white matter lesion in an MS brain. (**b**) Numerous FXR positive cells are detected within the center of the lesion (red box from panel A). (**c**) FXR positive cells are also detected at the edge of the lesion (blue box from panel A). (**d**) Staining for GPBAR1 within the white matter lesion revealed the presence of GPBAR1+ cells with an astrocytic morphology and also demonstrated positivity in vascular structures. (**e-f**) Double immunostaining for PLP (green) and GPBAR1 (red) shows the presence of GPBAR1+ cells (e) and vessels (f) in areas of demyelination. (**g-i**) Double immunostaining using GFAP (green) and GPBAR1 (red) demonstrates GPBAR1 staining in GFAP+ astrocytes in MS lesions. Scale bars: a, c=200um; b, d, e, f=100um; g, h, i=20um

### An endogenous bile acid blocks neurotoxic polarization of astrocytes and proinflammatory polarization of microglia

Since bile acid receptors are present on astrocytes and inflammatory cells in MS tissue, we investigated the effects of a bile acid – TUDCA on these cell populations *in vitro*. We chose TUDCA since it is currently being tested in multiple human trials in various neurological disorders including MS ^20^. We treated murine astrocytes with IL-1α, TNF-α, and C1q to polarize them to a neurotoxic (A1) phenotype^21^. We tested the effect of two doses of TUDCA and an agonist of GPBAR-1 (INT-777) in these polarizing conditions. TUDCA treatment blocked the upregulation of A1-specific gene transcripts in a dose dependent manner (Figure 4a). Interestingly, INT-777 partially recapitulated the effects of TUDCA on astrocyte polarization. Since ACM from A1 astrocytes can be toxic to neurons and oligodendrocytes, we tested the effects of ACM from non-polarized (A0), A1 polarized (vehicle) and A1 polarization in presence of TUDCA 70µM conditions, on the viability of murine oligodendrocytes (Figure 4b). As expected, ACM from TUDCA-treated astrocytes produced significantly less oligodendrocyte cell death compared to the vehicle condition in keeping with the changes noted in gene expression (Figure 4c and d).

**Fig. 4.**
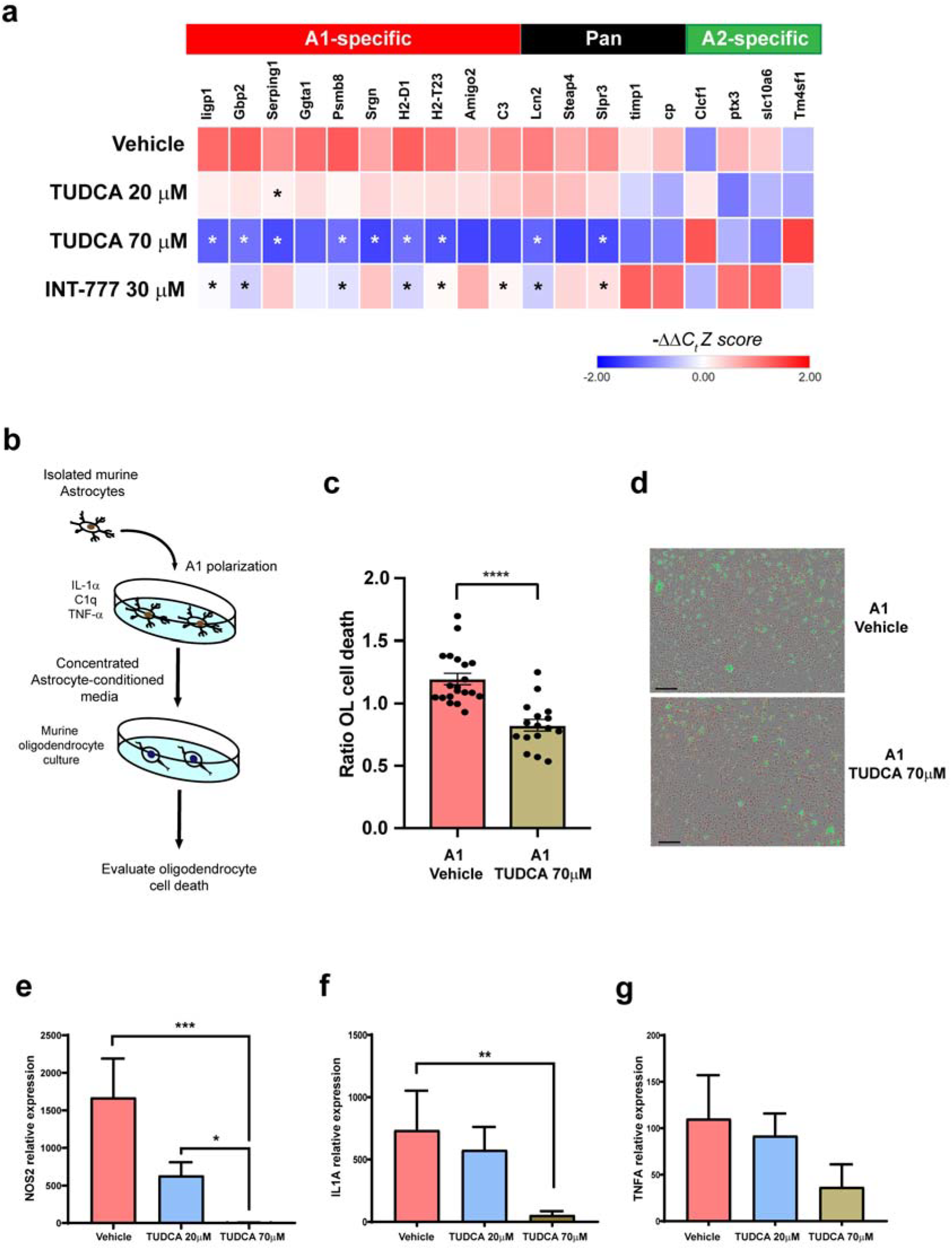
Endogenous bile acid blocks the neurotoxic polarization of astrocytes and pro-inflammatory polarization of microglia. **(a)** Astrocytes were isolated from neonatal mouse brains using ACSA2-bead selection and were polarized to an inflammatory phenotype using a cocktail of IL 1α, TNF-α and C1q for 24 hours either in the presence or absence of TUDCA or INT-777 (GPBAR1 agonist). Quantitative PCR of genes associated with various astrocyte phenotypes revealed that expression of multiple A1-specific genes was significantly down-regulated in the TUDCA 70µM condition compared to vehicle. We also noted down-regulation of several A1-specific genes with INT-777 treatment but to a lesser degree than TUDCA 70µM. Data are derived from three independent experiments with six biological replicates. Groups were compared with Kruskal-Wallis test with Dunn’s multiple comparisons test and asterisks represent p<0.05. **(b)** We concentrated astrocyte conditioned media from A0 (non-polarized), A1+ vehicle and A1+TUDCA 70µM condition and tested the effects on viability of murine oligodendrocytes over a 24-hour period. **(c)** We noted reduced oligodendrocyte cell death with TUDCA 70µM ACM compared to vehicle (normalized to cell death from A0 ACM). Data is derived from three independent biological replicates and groups were compared with an unpaired two-tailed Student’s t test. **(d)** Representative images depicting oligodendrocyte death testing ACM from the two conditions display greater cell death (Green) in vehicle condition compared to TUDCA 70µM. **(e)** Microglia were isolated from neonatal mouse brains using CD11b-bead selection and polarized to an inflammatory M1 phenotype by treatment with LPS and IFN-γ, either in the presence or absence of differing doses of TUDCA. We then isolated RNA to determine the expression of *NOS2* gene, which is a marker of M1 polarization and noted reduced expression in TUDCA treated conditions compared to vehicle. We also compared gene expression for *IL1A* **(f)** and *TNFA* **(g)**, which are factors produced by microglia critical for A1 polarization, in th various treatment conditions. We noted a significant reduction in gene expression for *IL1A* and a similar trend for *TNFA* in the TUDCA 70µM condition compared to vehicle. Data are derived from four independent experiments. We compared the groups with a one-way ANOVA with Dunnett’s test for multiple comparisons. In c and e-g bars represent mean with error bars represent s.e.m. (* p<0.05, ** p<0.01, *** p<0.005, **** p<0.001).

We polarized murine microglia to a pro-inflammatory phenotype by treating with IFN-γ and LPS, with or without TUDCA present. We noted a reduction in the expression of *NOS2* gene (Figure 4e) – a marker for M1 polarization and reduction in *IL1A* and *TNFA* transcription in the presence of TUDCA (Figure 4f and g). Thus, TUDCA, an endogenous bile acid, blocked the *in vitro* polarization of astrocytes to an A1 toxic phenotype and the polarization of microglia to a pro-inflammatory M1 phenotype.

### TUDCA supplementation ameliorates experimental autoimmune encephalomyelitis (EAE)

Since multiple circulating bile acid levels were lower in MS compared to healthy controls and TUDCA prevented the proinflammatory polarization of microglia and astrocytes, we next tested the effect of TUDCA supplementation in an animal model of MS - experimental autoimmune encephalomyelitis (EAE). EAE was induced by active immunization with MOG_35-55_ peptide and mice were randomized to receive either TUDCA 500 mg/kg orally or vehicle when they reached a disease severity score of 1 (Figure 5a). We noted reduction in the severity of EAE based on behavioral scores (Figure 5b) as well as pathological measures – reduced demyelination (Figure 5c and d), reduced infiltration of myeloid cells (Figure 5e and f) and reduced astrocytosis (Figure 5g and h). GPBAR1 receptors were noted on both astrocytes (GFAP+) and microglia/ macrophages (Mac-2+) cells in the spinal cords of mice with EAE (Supplementary Figure 4). We evaluated the presence of pro-inflammatory (M1) microglia/ macrophages using staining for iNOS and Iba-1 and noted a reduction in the proportion of iNOS+ Iba-1+ cells (Figure 5i and j). We also evaluated A1 astrocytes in EAE spinal cord tissue using staining for PSMB8 and GFAP, since PSMB8 was previously noted to be a marker of neurotoxic (A1) astrocytes^21^. A reduction in PSMB8+ GFAP+ cells was seen with TUDCA treatment (Figure 5k and l) suggesting a reduction in A1 astrocytes. These data are consistent with the *in vitro* data regarding the effects of TUDCA on both microglial and astrocytic polarization. Since CD4+ T cells are key players in the pathogenesis of the EAE model we tested the effects of TUDCA on CD4+ T cell proliferation and cytokine production *in vitro* and noted no direct anti-inflammatory effects of TUDCA on CD4+ T cells, as assessed by these measures (Supplementary Figure 5). These data suggest that TUDCA supplementation ameliorates neuroinflammation through reduced infiltration of immune cells and decreased microgliosis and astrocytosis.

**Fig. 5.**
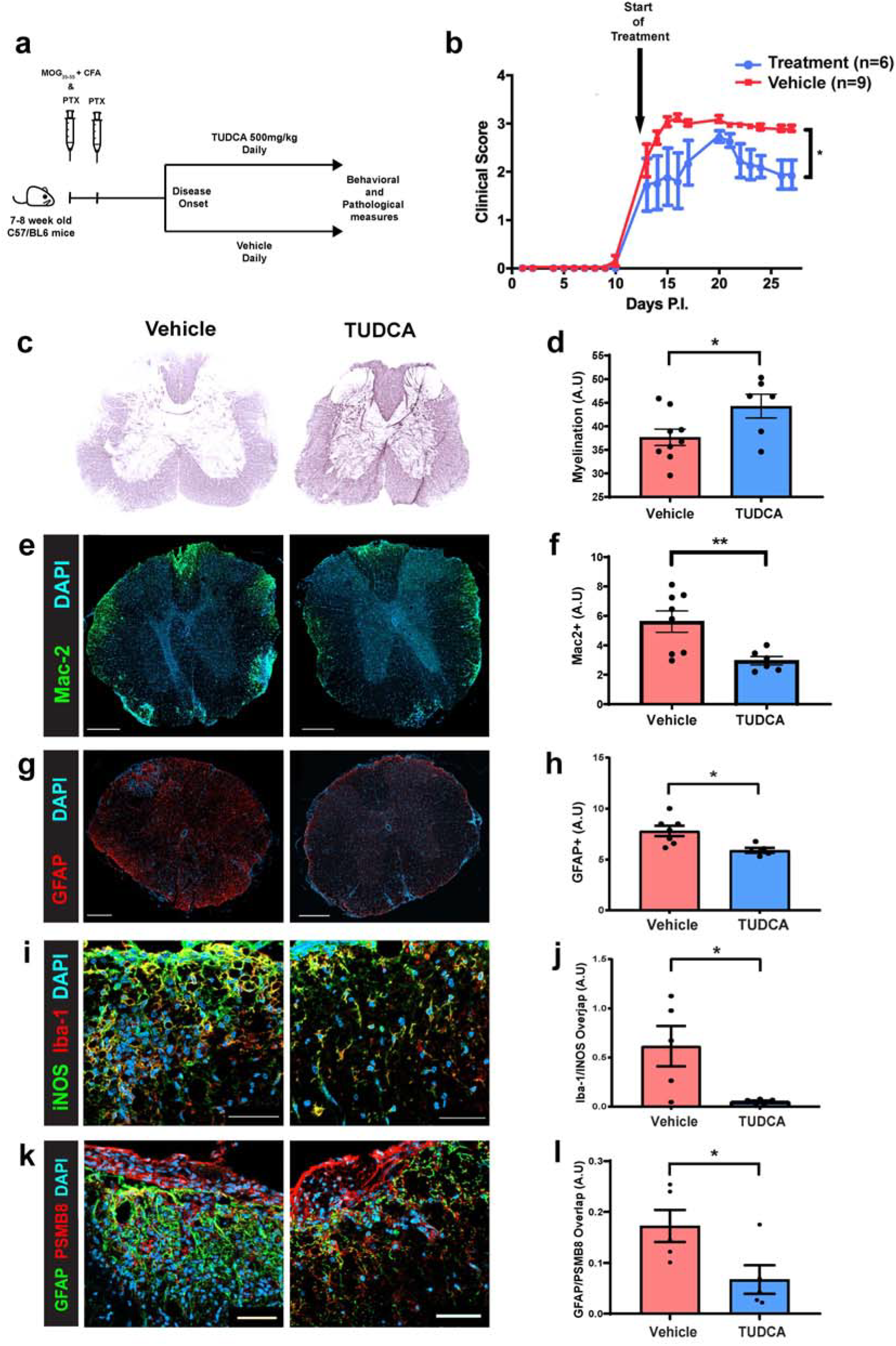
TUDCA supplementation ameliorates experimental autoimmune encephalomyelitis. **(a)** We immunized 7-8 week old female C57/BL6 mice subcutaneously with MOG_35-55_ and CFA in addition to intraperitoneal injection of PTX on day 0 and day 2. We then monitored these mice for signs of EAE and at disease onset we randomized mice to oral gavage with either TUDCA 500 mg/kg or vehicle till day 28 post-immunization. **(b)** Behavioral scores demonstrate that TUDCA treatment resulted in reduced disease severity of EAE (p value derived from a Mann-Whitney U test). **(c and d)** Representative images of Black-gold staining of spinal cords of mice with EAE treated with either TUDCA or vehicle demonstrate reduced demyelination with TUDCA treatment. **(e and f)** Immunostaining for infiltrating myeloid cells (Mac2+) demonstrated reduced infiltration in the TUDCA group compared to control. **(g and h)** Staining for GFAP demonstrates reduced astrocytosis in the TUDCA group compared to control. **(i and j** Staining for Iba-1 and INOS demonstrated decrease in INOS+ Iba-1+ cells (M1 macrophages/ microglia) in the TUDCA group compared to control. **(k and l)** We also noted reduced number of PSMB8+ GFAP+ cells (neurotoxic A1 astrocytes) in the TUDCA group compared to control. Scale bars: e and g = 200 µm, and k = 50 µm. Comparisons were made between the two treatment groups in d, f, h, j and l using an unpaired two-tailed Student’s t test. Bars in d, f, h, j and l represent mean with error bars representing s.e.m. (* p<0.05, ** p<0.005).

## Discussion

In this study, we demonstrate that bile acid metabolism is altered in both adult and pediatric-onset MS with lower levels of multiple circulating bile acid metabolites in the MS groups compared to healthy controls. We also demonstrate the presence of two key bile acid receptors – FXR and GPBAR1 - on glial and immune cells in MS brain tissue. We then identify the direct anti-inflammatory effects of an endogenous bile acid – TUDCA on astrocytes and microglia *in vitro* and show that supplementation with TUDCA ameliorates disease in a mouse model of MS.

Previous studies have demonstrated that bile acids have anti-inflammatory effects on myeloid cells, which include the prevention of M1 polarization and blockade of the NLRP3 inflammasome pathway^16^. Cues from microglia influence astrocytic phenotype and impact neuroinflammation^21,22^. We confirmed that TUDCA not only prevented pro-inflammatory polarization of microglia, but also reduced production of factors implicated in the polarization of astrocytes to an A1 phenotype. Besides the potential indirect effect through prevention of microglial polarization, we identified that TUDCA can act directly on astrocytes and block A1 polarization. TUDCA treatment decreased upregulation of A1-specific genes and the change in gene expression resulted in a functional change - reduced toxicity of ACM to oligodendrocytes. To the best of our knowledge, this is the first example of an endogenous metabolite that directly blocks the polarization of astrocytes to a toxic phenotype. This has important implications since this subset of astrocytes could play a critical role in neurodegeneration in disorders such as Parkinson’s disease (PD), AD and MS^21^. Since, we noted that astrocytes in both humans and mice express the GPBAR1 receptor, we tested whether the effects of TUDCA on astrocyte polarization could be replicated using a GPBAR1 agonist – INT777. We identified partial replication of the effects of TUDCA with INT777 suggesting that this beneficial effect may be partially mediated through GPBAR1. A previous study has also reported that the GPBAR1 receptor may mediate anti-inflammatory effects of TUDCA on microglial cells^16^. Thus, while our data suggests that GPBAR1 may be important for the effects of bile acids on astrocyte polarization, the precise mechanism requires further study.

The identification of altered serum levels of bile acids in MS is not entirely unexpected given that gut dysbiosis has been reported in MS patients in several studies^9,23–25^. Since modification of bile acids by the gut microbiota is a key step in their metabolism, the alteration in abundance of bacterial species that are involved in these processes could potentially explain altered circulating levels of bile acid metabolites in MS. Indeed, bacterial species including *Clostridium* and *Parabacteroides* which have been noted to be reduced in abundance in MS are important players in bile acid metabolism and have been linked to alteration in circulating levels of certain bile acids^10,19,26^. The change in ratios of secondary to primary bile acids in both synthesis pathways also points to the potential role of gut dysbiosis as a key factor driving this change. The observation that these changes was more marked in progressive MS parallels the trend noted in AD, where the change in bile acid profiles was more marked in AD than in mild cognitive impairment (MCI)^4^. This potentially suggests a relationship between greater abnormality in bile acid metabolism and neurodegeneration.

As the makeup of the gut microbiota is linked to an individual’s genome, circulating metabolite levels could also be driven by specific genetic factors^26^. In a recent study, vitamin D receptor (VDR) polymorphisms were demonstrated to alter *Parabacteroides* abundance and ultimately impact the level of specific bile acids^26^. Similarly, other MS-related genetic risk alleles could alter the gut microbiota and ultimately lead to metabolic changes such as those noted in our study. For example, *ZFP36L1* (a gene identified in several MS GWAS studies) was recently demonstrated to modulate bile acid biosynthesis^27^.

Besides altered gut microbial bile acid metabolism, altered bile acid profiles could result from abnormal endogenous production. Since cholesterol is the precursor of bile acids, alterations in cholesterol metabolism could potentially underlie changes in bile acid abundance. Cholesterol metabolism is intricately linked to T cell and innate immune cell function^28^. Inflammation can alter cholesterol metabolism in the CNS and this was the major abnormal pathway in a recent study evaluating spinal cord astrocyte gene expression profiles in EAE^11^. These data suggest that inflammation can alter endogenous cholesterol metabolism, in the CNS and the periphery and could be a factor underlying the altered bile acid levels in MS patient circulation.

We examined the distribution of two key bile acid receptors in MS brains. FXR is a nuclear bile acid receptor present in MS white matter lesions from RRMS patients – primarily in immune cells^29^. We found that in addition to active lesions, FXR expression can also be found in chronic MS lesions. We also identified the presence of GPBAR1 on astrocytes and vascular structures in MS lesions. The expression of GPBAR1 on astrocytes raises the possibility that the circulating bile acid metabolites could potentially alter neuroinflammation through effects on these cells. In animal models, GPBAR1 expressed on the endothelium helped regulate chemotaxis of immune cells^30^ and the presence of this receptor on vessels in the MS brain raises the possibility that it could help regulate entry of immune cells into the CNS parenchyma. Overall the presence of both nuclear and surface receptors in MS brains reinforce the relevance of altered circulating bile acid levels in MS.

Previous studies have demonstrated that bile acid levels are reduced in chronic EAE, providing a strong rationale for testing bile acid supplementation in this animal model^12^. Obeticholic acid - a synthetic bile acid and an agonist for FXR led to amelioration of EAE^14^. Previous work also established that FXR positive cells are present in the CNS in EAE and FXR agonists can lead to amelioration of the disease^13^. However, despite the expression of FXR in glial cells, a previous study demonstrated that an FXR agonist did not result in anti-inflammatory effects in astrocytes or microglia suggesting that the beneficial effect of FXR agonism in EAE is mediated through effects on peripheral myeloid cells^31^. We demonstrated that astrocytes and myeloid cells in the CNS of EAE mice also express GPBAR1, thus providing additional support for evaluating the *in vivo* effects of TUDCA supplementation in this model. Besides identifying a beneficial effect of TUDCA supplementation on EAE severity, we also noted reduced numbers of iNOS-expressing microglia/ macrophages (M1 phenotype) and PSMB8-expressing astrocytes (A1 phenotype). This provides *in vivo* corroboration of the anti-inflammatory effects of bile acid supplementation that we noted initially in our *in vitro* experiments. The primary mechanism of the beneficial effects of bile acid supplementation in the EAE model – whether related to GPBAR1 activation in glial cells or FXR activation in peripheral myeloid cells requires further study.

Our study has several strengths. We demonstrated abnormal bile acid metabolism in two cohorts from different centers and in different age groups. Besides evaluating this observation in humans, we also evaluated the effects of bile acids *in vitro* and in an animal model of MS. The limitations of the current study include the cross-sectional nature of the metabolomics analyses precluding an assessment of how bile acids change over time in MS and whether they are affected by specific MS disease-modifying therapies. Fewer bile acid metabolites were measured in the targeted cohort compared to the untargeted panel, i.e. not all metabolites were present in both data sets for direct comparison. However, we did note that the majority of metabolites demonstrated consistent changes across all the cohorts. Finally, human samples were obtained after disease onset which precludes evaluation of causality.

Future studies will help further define the relationship between bile acid levels and MS disease course. Additionally, ongoing clinical trials of supplementation with TUDCA in MS (NCT03423121) are likely to provide us with information regarding the effects of this intervention on circulating bile acid levels and ultimately on MS activity and disease course.

The results of this study expand our understanding of how alterations in the metabolome may affect aspects of MS disease pathophysiology and potentially impact processes that mediate neurodegeneration in multiple neurological diseases. We also provide a paradigm for future studies to identify additional metabolic pathways that could be targeted to modify the course of MS.

## Methods

### Patient cohorts

We recruited adult MS patients and healthy controls from the Johns Hopkins MS Center. Participants underwent phlebotomy for blood collection in addition to collection of clinical and demographic data. Blood was processed using our standard protocol and plasma stored at −80 C until the time of metabolomics analyses. We had two adult cohorts in this study – a discovery cohort consisting of 56 RRMS, 51 PMS and 52 HC that underwent global metabolomics analysis and a validation cohort which consisted of 75 HC, 50 RRMS and 125 PMS that underwent targeted metabolomics analysis. Pediatric MS patients and healthy controls (n=31 each) were recruited from the UCSF Pediatric MS clinic. Participants underwent phlebotomy and blood was processed and serum stored at −80 C until the time of metabolomics analyses ^32^.

### Global Metabolomics analyses

Global metabolomics analyses were performed at Metabolon Inc (Durham, NC) as previously described ^33^. In brief, recovery standards were added prior to the extraction process for quality control. To remove protein and to recover chemically diverse metabolites, proteins were precipitated with methanol under vigorous shaking (Glen Mills Genogrinder 2000) for 2 minutes, followed by centrifugation. The resulting extract was divided into 5 fractions: analysis by ultra-high-performance liquid chromatography–tandem mass spectrometry (UPLC-MS/MS; positive ionization), UPLC-MS/MS (negative ionization), UPLC-MS/MS polar platform (negative ionization), gas chromatography–mass spectrometry, and one aliquot was reserved for backup. Metabolite identification was performed by automated comparison of the ion features in the study samples to a reference library of standard metabolites. Quantification of peaks was performed using area under the curve. Raw values for the area counts, for each metabolite were normalized (correcting for variation resulting from instrument inter-day tuning differences) by the median value for each run day.

### Targeted metabolomics assay

The targeted bile acid panel measured all of the major human primary and secondary bile acids and their respective glycine and taurine conjugates: Cholic Acid (CA), Chenodeoxycholic Acid (CDCA), Deoxycholic Acid (DCA), Lithocholic Acid (LCA), Ursodeoxycholic Acid (UDCA), Glycocholic Acid (GCA), Glycochenodeoxycholic Acid (GCDCA), Glycodeoxycholic Acid (GDCA), Glycoursodeoxycholic Acid (GUDCA), Taurocholic Acid (TCA), Taurochenodeoxycholic Acid (TCDCA), Taurodeoxycholic Acid (TDCA), Taurolithocholic Acid (TLCA), Tauroursodeoxycholic Acid (TUDCA), and Glycolithocholic Acid (GLCA). Bile acid concentrations were analyzed by LC-MS/MS. The individual analyte quantitation ranges (based on the analysis of 50.0 µl plasma) are listed in Supplementary Table 2.

Calibration samples were prepared at eight different concentration levels by spiking a phosphate buffered saline/bovine serum albumin (PBS/BSA) solution with corresponding calibration spiking solutions. Calibration samples, study samples, and quality control samples were spiked with a solution of labeled internal standards and subjected to protein precipitation with an organic solvent (acidified MeOH). Following centrifugation, an aliquot of the organic supernatant was evaporated to dryness in a gentle stream of nitrogen. The dried extracts were reconstituted and injected onto an Agilent 1290/Sciex QTrap 6500 LC-MS/MS or Agilent Infinity II/Sciex TripleQuad 6500+ LC-MS/MS system equipped with a C18 reverse phase HPLC column. The mass spectrometer was operated in negative mode using electrospray ionization. The peak area of each bile acid parent (pseudo-MRM mode) or product ion was measured against the peak area of the respective internal standard parent (pseudo-MRM mode) or product ion. Quantitation was performed using a weighted linear least squares regression analysis generated from fortified calibration standards prepared immediately prior to each run.

Single level pooled QC samples were used with most bile acid concentrations at the endogenous level. Due to low abundance of TUDCA, TCA, TLCA, GLCA, and LCA in this lot, the QCs were up-spiked to typically observed levels (standard C to D range). Due to multiplexing 15 analytes, only a single concentration level of QC samples was used. Raw data were collected and processed using AB SCIEX software Analyst 1.6.3. Data reduction was performed using Microsoft Excel 2013.

Sample analysis was carried out in the 96-well plate format containing two calibration curves and six QC samples (per plate) to monitor method performance. Four sample batches were prepared and analyzed. Following analysis, four samples were found to be above the limit of quantitation (ALOQ) and therefore re-analyzed at a 5-fold dilution. Precision was evaluated using the corresponding QC replicates in each sample run. Intra-run and inter-run precision (%CV) of all analytes met acceptance criteria.

### Immunohistochemistry of MS autopsy tissue

All brains were collected as part of the tissue procurement program approved by the Cleveland Clinic Institutional Review Board. Multiple sclerosis (MS) brain tissues were characterized for demyelination by immunostaining using 30 µm fixed tissue sections and proteolipid protein (PLP) using protocols described previously ^34–36^. Briefly, fixed blocks of the tissue were cut on a sliding microtome, microwaved in 10 mM citric acid buffer (pH 6.0) and immunostained by the avidin-biotin complex procedure with diaminobenzidine (DAB) using rat anti-proteolipid protein (1:250, gift from Wendy Macklin, University of Colorado, Denver), mouse anti-FXR (1:500, Perseus Proteomics, Tokyo, Japan), rabbit anti-TGR5 (1:250, Thermo Fisher Scientific, Rockford, IL) and anti-GFAP (1:1000, Millipore, Burlington, MA) as described previously (refs). Adjacent sections were used for double labeling (PLP-TGR5 and GFAP-TGR5) using respective secondary antibodies (biotinylated donkey anti-rat IgG, donkey anti-mouse IgG, and donkey anti-rabbit IgG (1:500, Vector Laboratories, Burlingame, CA), and Alexa-488 donkey anti-rat IgG, Alexa-488 donkey anti-mouse IgG and Alexa-594 donkey anti-rabbit IgG (1:500, Invitrogen, Carlsbad, CA). Immunofluorescent-labeled tissues were analyzed using a Leica DM5500 upright microscope (Leica Microsystems, Exton, PA). PLP-TGR5 and GFAP-TGR5 images were collected from corresponding sections. Resultant images were processed using Fiji version (http://fiji.sc) of the free image processing software ImageJ (NIH, http://rsbweb.nih. gov/ij) as described before ^34–36^.

### Murine Microglia Primary Cultures

Microglia were isolated from brains of post-natal day 3 – 5 mouse pups. After dissecting off the meninges, whole brains underwent mechanical and enzymatic dissociation using Neural Tissue Dissociation Kit - Papain (Miltenyi, Bergisch Gladbach, Germany). Microglia were then positively selected using CD11b magnetic beads (Miltenyi, Bergisch Gladbach, Germany) and seeded onto poly-L-lysine-coated, 6-well plates at a density of 3 × 10^5^ cells/mL in DMEM/F12 containing 10% fetal bovine serum, 1% penicillin/streptomycin, and 1% Glutamax. Microglial purity was confirmed by flow cytometry (CD11b+ CD45-low ACSA2-A2B5-O4-). After 24 h, cells were left unstimulated or stimulated with recombinant murine IL-4 (25 ng/mL, PeproTech, Rocky Hill, NJ) or a combination of lipopolysaccharide (LPS, 100 ng/mL) and recombinant murine IFN□ (25 ng/mL, PeproTech, Rocky Hill, NJ). Concurrently, cells were treated with various bile acid conditions: 20 µM TUDCA and 70 µM TUDCA. After 18 h, supernatants were collected, and cells were lysed and prepared for qPCR analysis.

### Murine Astrocyte Primary Cultures

Astrocytes were isolated from brains of post-natal day 3 – 5 mouse pups. Brains were enzymatically dissociated similarly to microglia (see above). Cells were Fc blocked with anti-CD16/CD32 antibodies then positively selected using ACSA-2 magnetic beads (Miltenyi, Bergisch Gladbach, Germany) and seeded onto poly-L-lysine-coated, 6-well plates at a density of 2.5 × 10^5^ cells/mL. Astrocytes were cultured in serum-free media containing 50% DMEM, 50% neurobasal, 1% penicillin/streptomycin, 1% glutamax, 1 mM sodium pyruvate, N-acetyl cysteine (5 μg/mL), 1 × SATO, and supplemented with HB-EGF (5 ng/mL, PeproTech, Rocky Hill, NJ), as previously described ^21^. Astroglial purity was confirmed by flow cytometry (ACSA2+ CD11b-CD45- A2B5- O4-). Astrocytes were grown until cultures reached a desired density (approximately 7 days) with media changes every 2 days. Cells were then left unstimulated or stimulated to generate A1 polarized astrocytes by adding recombinant rat Il-1α (3 ng/mL, PeproTech, Rocky Hill, NJ), recombinant human TNFα (30 ng/mL, Sigma), and C1q (400 ng/mL, MyBioSource, San Diego, CA). Concurrently, cells were treated with various conditions: 20 μM TUDCA, 70 μM TUDCA and 30 μM INT777. After 24 h, supernatants were collected and cells were lysed and prepared for qPCR analysis.

### Cell Viability Assay

Cells were seeded onto 96-well plates (50,000 microglia or astrocyte/well) and grown for 7 days (astrocyte) or 24 h (microglia). To assess toxicity of bile acid treatment or other treatments, dead cells were stained with Cytotox Green or Cytotox Red Reagent (IncuCyte, Ann Arbor, MI) following the manufacturer’s instructions. Plates were monitored and images were captured with the IncuCyte S3 every 2 hours over a period of 18 hours (microglia) or 24 hours (astrocytes). Two experiments with six replicates of each condition per experiment were quantified. For each well 9 images were acquired, and fluorescence positive cells were quantified using the IncuCyte S3 software.

### Quantitative PCR

RNA was isolated from vehicle- or bile acid-treated microglia and astrocyte cultures using RNeasy Plus Mini Kit (Qiagen, Hilden, Germany). First-strand cDNA was synthesized by reverse transcription using the iScript cDNA Synthesis Kit (Bio-Rad, Hercules, CA). Samples were quantified and amplified using SensiMix (Bioline, London, UK), a SYBR based reagent and a CFX384 Touch Real-Time PCR Detection System (Bio-Rad, Hercules, CA). Primer sequences (IDT, San Jose, CA) are listed in Supplementary Table 7. The threshold cycle (CT) values for the target genes were normalized to b-actin reference gene (ΔCT) and experimental controls (ΔΔCT).

### Assessment of oligodendrocyte killing by A1 astrocytes

Astrocyte conditioned media was collected with complete Mini-EDTA free protease inhibitor cocktail (Sigma, St. Louis, MO) and concentrated with Amicon Ultra-15 30kDa filters (e, Burlington, MA) by centrifuging 4000g for 20 minutes. Protein concentration was measured with Bradford assay.

Oligodendrocytes were isolated by sequential immunopanning. P6-P8 mouse pup cerebral cortices were enzymatically and mechanically dissociated with papain (MACS Miltenyi Biotec, Bergisch Gladbach, Germany) to generate a single cell suspension. Cell suspension was sequentially plated on series of negative selection antibody-coated plates to remove endothelial cells (BSL1, Vector L1100), microglia (CD11b clone OX-42, Biorad, Hercules, CA) then positive-selection plate for oligodendrocyte progenitors (PDGFR, BD Biosciences, San Jose, CA). Isolated oligodendrocyte progenitors were plated on poly-D-lysine coated coverslips in 24 well plates in oligodendrocyte base media (high glucose DMEM, 100U/ml penicillin, 100µg/ml streptomycin, 1mM sodium pyruvate, 4mM L-glutamine, 5ug/ml N-acetyl cysteine, 1× SATO, 1× B27, 5µg/ml insulin, 10ng/ml d-biotin, 1× trace elements B, 4.2µg/ml forskolin) supplemented with 20ng/ml PDGFAA (Peprotech, Rocky Hill, NJ). Oligodendrocyte progenitors were allowed to proliferate and recover for several days and media was replaced with differentiation media- oligodendrocyte base media with 40ng/ml T3. After 24 hours of differentiation, 50µg/ml of concentrated astrocyte conditioned media was added to differentiated oligodendrocytes with Cytotox green (IncuCyte, Ann Arbor, MI) and NucLight rapid red (IncuCyte, Ann Arbor, MI). Plates were monitored in IncuCyte S3 Live-Cell Analysis system incubator and 10× images were captured. IncuCyte S3 program was used to set fixed threshold masks to quantify Cytotox and rapid red nuclear labeling. Images that met defined density thresholds of 5-50% on phase contrast and 1-25% on rapid red nuclear labelling were quantified. Five experimental replicate sets of astrocyte conditioned media (A0, A1 and A1+ TUDCA 70µM treated) with 3-4 wells per replicate were quantified. Data was analyzed in GraphPad Prism 8. For each well 3-14 images were quantified to calculate the mean and SEM of ratio of cytotox confluence/rapid red confluence per well. The mean of A0 cell death ratio was set to baseline of 1 and A1 and A1 + TUDCA 70µM conditions were compared to A0 baseline. Two-way ANOVA was used to compare means between well replicates and experimental samples. Unpaired Welch’s t test was used to compare two sample group means.

### Experimental Autoimmune Encephalomyelitis induction and treatment

C57/BL6J mice were purchased from Jackson Laboratories. All mice were maintained in a federally approved animal facility at Johns Hopkins University in accordance with Institutional Animal Care and Use Committee. Female mice of 7-8 weeks of age were used in these experiments.

EAE was induced by s.c injection over two sites in the flank with 100 µg myelin oligodendrocyte glycoprotein (MOG_35-55_) peptide (Johns Hopkins Peptide Synthesis Core Facility) emulsified in Complete Freund’s Adjuvant (Difco, Detroit, MI). A total of 250 ng of pertussis toxin (List Biologicals, Campbell, CA) was administered by i.p. injection at the time of immunization and 48 hours later. Mice were monitored for signs of EAE and scored based on the following criteria: 0 – no disease, 1 – limp tail, 2 – hind limb weakness, 3 – hind limb paralysis, 4 – forelimb weakness in addition to hind limb paralysis and 5 – death due to EAE.

At the onset of clinical disease mice were randomized to daily treatment with either TUDCA 500 mg/kg (dissolved in PBS) or vehicle (PBS) administered by oral gavage. Mice were treated till day 28 post immunization. Scoring of disease severity was performed by raters blinded to treatment allocation.

### Immunohistochemistry of EAE tissue

EAE mice were deeply anesthetized, then perfused transcardially with chilled PBS followed by 4% PFA. Spinal columns were dissected and post-fixed in 4% PFA overnight followed by cryoprotection with 30% sucrose for 48 hours. The spinal cords were then dissected and embedded in OCT before freezing. Tissues were sectioned on a cryostat (10 um sections) and mounted on glass slides (Super-frost Plus; Fisher, Hampton, NH). Blocking and permeabilization of sections was performed in PBS containing 5% normal goat serum (NGS) and 0.4% Triton X-100 for 1 h at room temperature followed by incubation overnight at 4°C in PBS containing 3% normal goat serum, 0.1% Triton X-100, and primary antibody (Supplementary Table 8). Sections were incubated in secondary antibodies conjugated to Alexa fluorophores (1:1000, Invitrogen, Carlsbad, CA) for 1 h at room temperature before mounting in anti-fade reagent with DAPI (Prolong Gold Anti-fade, Gibco, Gaithersburg, MD).

Black Gold staining (Black Gold II myelin staining kit, Millipore, Burlington, MA) was performed according to the manufacturer’s instructions. In short, slides were incubated in 0.3% Black Gold at 60°C for 12–20 min until the thinnest fibers were stained. The slides were then fixed in 1% sodium thiosulfate for 3 min, counterstained with 0.1% Cresyl Violet (Millipore, Burlington, MA) for 3 min, dehydrated using a series of gradated alcohols, cleared in xylene, and cover slipped with VectaMount Permanent Mounting Media (Vector Laboratories, Burlingame, CA).

Slides were imaged alternatively on a Keyence BZ-X710 All-in-One Fluorescence Microscope (Bright-field; 20×), Zeiss Axio Observer Z1 epifluorescence microscope (20×), and Zeiss LSM 800 confocal microscope (Apochromat 20×/0.8; 516px × 516px). We imaged 5-6 sections per mouse in 5-8 mice in each group. National Institutes of Health ImageJ software was used to create a binary image of the staining and to subsequently quantify staining intensity and area.

### Murine T-Cell Proliferation Study

8-12 week old C57/BL6J mice (Jackson Laboratory, Bar Habor, MA) were euthanized with isoflurane followed by cervical dislocation. Spleen and inguinal lymph nodes were collected and dissociated by passing through 100µm cell strainers. Red blood cells were osmotically lysed and CD4+ T-cells were isolated using a MojoSort Mouse CD4+ T Cell Isolation Kit (Biolegend, San Diego, CA). Purified CD4s were labeled with Cell Proliferation Dye eFluor450 (ThermoFisher, Waltham, MA) then cultured with plate-bound anti-CD3 and soluble anti-CD28 (BD Biosciences, San Jose, CA) at 2µg/ml in complete RPMI. After 3 days cells were collected and re-activated with Cell Stimulation Cocktail + Protein Transport Inhibitor (ThermoFisher, Waltham, MA) for 5 hours then washed, stained for viability with Live/Dead Aqua (ThermoFisher, Waltham, MA), stained for surface markers, fixed and permeabilized with IC Fix buffer followed by Permeabilization Buffer (ThermoFisher, Waltham, MA) then stained for cytokines in permeabilization buffer. Cells were washed twice and run on a MACSQuant 10 cytometer (Miltenyi, San Diego, CA). Data was analyzed with FlowJo V10 software (BD Biosciences, San Jose, CA). We performed three independent experiments with a total of five biological replicates. All kits and reagents were used following manufacturers recommended protocols.

### Statistical Analysis

For untargeted and targeted metabolomics data we excluded any metabolites that had greater than 30% missing values and then normalized and scaled the metabolite relative abundance. The pre-processed data was then used to generate pathway deregulation scores with the *Pathifier* package in R ^37^. The pathway deregulation scores indicate the distance of an individual from a principal components curve generated using data from control subjects. The scores range from 0 to 1 with higher scores indicating greater abnormality.

The individual metabolite abundances and the pathway deregulation scores were then compared between groups using multivariate linear regression models adjusting for age, sex and race. We also generated heatmaps of Z-scores of the relative abundances of each of the bile acid metabolites using Morpheus (https://software.broadinstitute.org/morpheus/). We utilized Stata version 15 for regression analyses and Graphpad Prism 8.0 to generate all graphs.

The -ΔΔCT values for astrocyte gene expression were compared between culture conditions using Kruskal-Wallis test with Dunn’s multiple comparisons test. Heatmap for gene expression profiles of astrocyte culture were also created using Morpheus (https://software.broadinstitute.org/morpheus/). Gene expression values from microglial PCR data were compared using one-way ANOVA between groups with Dunnett’s correction for multiple comparisons.

We compared EAE scores between TUDCA and vehicle groups using a two-tailed Mann-Whitney U test. We compared quantification of staining for various markers between the groups using an unpaired two-tailed Student’s t test.

### Study Approvals

All participants at both centers provided informed consent prior to enrollment and the studies were approved by the Institutional Review Board at each site. For all animal experiments we adhered to the guidelines in the NIH Guide for the Care and Use of Laboratory Animals. Protocols were approved by the Institutional Animal Care and Use Committee.

## Author contributions

PB was involved in conceptualization, data curation, formal analysis, funding acquisition, investigation, supervision, visualization and writing – original draft, LM was involved in data curation, investigation, validation, writing – review & editing, MDS was involved in investigation, visualization and writing – review & editing, EH was involved in investigation, visualization, writing – review & editing, KCF was involved in investigation, formal analysis, writing – review & editing, KM was involved in investigation, visualization, writing – review & editing, SK was involved in investigation, visualization, writing – review & editing, AR was involved in investigation, visualization, writing – review & editing, JGC was involved in investigation, visualization, writing – review & editing, SS was involved in investigation, visualization, writing – review & editing, KV was involved in investigation, visualization, writing – review & editing, CV was involved in investigation, visualization, writing – review & editing, ESS was involved in investigation, visualization, writing – review & editing, BN was involved in investigation, validation, writing – review & editing, RD was involved in investigation, visualization, writing – review & editing, EMM was involved in investigation, supervision, writing – review & editing, EW PAC was involved in conceptualization, methodology, funding acquisition, supervision and writing – review & editing.

## Acknowledgments

PB was supported by a John F Kurtzke Clinician Scientist Development Award from the American Academy of Neurology and a Career Transition Award (TA-1503-03465) from the National Multiple Sclerosis Society, RD and CV are supported by NINDS (NS096148). This study was partly supported by a Marilyn Hilton Award for Innovation in MS research to PAC and a research grant from the National Multiple Sclerosis Society (RG-1707-28601).

## Data Availability Statement

Metabolomics data will be deposited to NIH Metabolomics Workbench.

## References

1. Ridlon, J. M., Kang, D. J., Hylemon, P. B. & Bajaj, J. S. Bile acids and the gut microbiome. Curr. Opin. Gastroenterol. 30, 332–338 (2014).

2. McMillin, M. & DeMorrow, S. Effects of bile acids on neurological function and disease. FASEB J. 30, 3658–3668 (2016).

3. Guo, C. et al. Bile Acids Control Inflammation and Metabolic Disorder through Inhibition of NLRP3 Inflammasome. Immunity 45, 802–816 (2016).

4. MahmoudianDehkordi, S. et al. Altered bile acid profile associates with cognitive impairment in Alzheimer’s disease-An emerging role for gut microbiome. Alzheimers. Dement. 0, (2018).

5. Isobe, N. et al. An ImmunoChip study of multiple sclerosis risk in African Americans. Brain 138, 1518–30 (2015).

6. The International Multiple Sclerosis Genetics Consortium (IMSGC). Evidence for Polygenic Susceptibility to Multiple Sclerosis-The Shape of Things to Come. Am. J. Hum. Genet. 86, 621–625 (2010).

7. Ascherio, A. et al. Vitamin D as an Early Predictor of Multiple Sclerosis Activity and Progression. JAMA Neurol. (2014). doi:10.1001/jamaneurol.2013.5993

8. Bhargava, P. & Mowry, E. M. Gut microbiome and multiple sclerosis. Curr. Neurol. Neurosci. Rep. 14, 492 (2014).

9. Jangi, S. et al. Alterations of the human gut microbiome in multiple sclerosis. Nat. Commun. 7, 12015 (2016).

10. Cekanaviciute, E. et al. Gut bacteria from multiple sclerosis patients modulate human T cells and exacerbate symptoms in mouse models. Proc. Natl. Acad. Sci. U. S. A. 114, 10713–10718 (2017).

11. Itoh, N. et al. Cell-specific and region-specific transcriptomics in the multiple sclerosis model: Focus on astrocytes. Proc. Natl. Acad. Sci. 115, E302–E309 (2018).

12. Mangalam, A. et al. Profile of Circulatory Metabolites in a Relapsing-remitting Animal Model of Multiple Sclerosis using Global Metabolomics. J. Clin. Cell. Immunol. 4, (2013).

13. Hucke, S. et al. The farnesoid-X-receptor in myeloid cells controls CNS autoimmunity in an IL-10-dependent fashion. Acta Neuropathol. 132, 413–31 (2016).

14. Ho, P. P. & Steinman, L. Obeticholic acid, a synthetic bile acid agonist of the farnesoid X receptor, attenuates experimental autoimmune encephalomyelitis. Proc. Natl. Acad. Sci. U. S. A. 113, 1600–5 (2016).

15. Lewis, N. D. et al. A GPBAR1 (TGR5) small molecule agonist shows specific inhibitory effects on myeloid cell activation in vitro and reduces experimental autoimmune encephalitis (EAE) in vivo. PLoS One 9, e100883 (2014).

16. Yanguas-Casás, N., Barreda-Manso, M. A., Nieto-Sampedro, M. & Romero-Ramírez, L. TUDCA: An Agonist of the Bile Acid Receptor GPBAR1/TGR5 with Anti-Inflammatory Effects in Microglial Cells. J. Cell. Physiol. (2016). doi:10.1002/jcp.25742

17. Gaspar, J. M. et al. Tauroursodeoxycholic acid protects retinal neural cells from cell death induced by prolonged exposure to elevated glucose. Neuroscience 253, 380–388 (2013).

18. Mortiboys, H. et al. UDCA exerts beneficial effect on mitochondrial dysfunction in LRRK2(G2019S) carriers and in vivo. Neurology 85, 846–52 (2015).

19. Staley, C., Weingarden, A. R., Khoruts, A. & Sadowsky, M. J. Interaction of gut microbiota with bile acid metabolism and its influence on disease states. Appl. Microbiol. Biotechnol. 101, 47–64 (2017).

20. Elia, A. E. et al. Tauroursodeoxycholic acid in the treatment of patients with amyotrophic lateral sclerosis. Eur. J. Neurol. (2015). doi:10.1111/ene.12664

21. Liddelow, S. A. et al. Neurotoxic reactive astrocytes are induced by activated microglia. Nature 541, 481–487 (2017).

22. Rothhammer, V. et al. Microglial control of astrocytes in response to microbial metabolites. Nature 557, 724–728 (2018).

23. Cantarel, B. L. et al. Gut microbiota in multiple sclerosis: possible influence of immunomodulators. J. Investig. Med. 63, 729–34 (2015).

24. Tremlett, H. et al. Gut microbiota in early pediatric multiple sclerosis: a case-control study. Eur. J. Neurol. 23, 1308–1321 (2016).

25. Chen, J. et al. Multiple sclerosis patients have a distinct gut microbiota compared to healthy controls. Sci. Rep. 6, 28484 (2016).

26. Wang, J. et al. Genome-wide association analysis identifies variation in vitamin D receptor and other host factors influencing the gut microbiota. Nat. Genet. 48, 1396–1406 (2016).

27. Tarling, E. J. et al. RNA-binding protein ZFP36L1 maintains posttranscriptional regulation of bile acid metabolism. J. Clin. Invest. 127, 3741–3754 (2017).

28. Puleston, D. J., Villa, M. & Pearce, E. L. Ancillary Activity: Beyond Core Metabolism in Immune Cells. Cell Metab. 26, 131–141 (2017).

29. Hucke, S. et al. The farnesoid-X-receptor in myeloid cells controls CNS autoimmunity in an IL-10-dependent fashion. Acta Neuropathol. 132, 413–431 (2016).

30. Kida, T., Tsubosaka, Y., Hori, M., Ozaki, H. & Murata, T. Bile Acid Receptor TGR5 Agonism Induces NO Production and Reduces Monocyte Adhesion in Vascular Endothelial Cells. Arterioscler. Thromb. Vasc. Biol. 33, 1663–1669 (2013).

31. Albrecht, S. et al. Activation of FXR pathway does not alter glial cell function. J. Neuroinflammation 14, 66 (2017).

32. Nourbakhsh, B. et al. Altered tryptophan metabolism is associated with pediatric multiple sclerosis risk and course. Ann. Clin. Transl. Neurol. (2018). doi:10.1002/acn3.637

33. Bhargava, P., Fitzgerald, K. C., Calabresi, P. A. & Mowry, E. M. Metabolic alterations in multiple sclerosis and the impact of vitamin D supplementation. JCI Insight 2, (2017).

34. Trapp, B. D. et al. Axonal transection in the lesions of multiple sclerosis. N. Engl. J. Med. 338, 278–85 (1998).

35. Dutta, R. et al. Mitochondrial dysfunction as a cause of axonal degeneration in multiple sclerosis patients. Ann. Neurol. 59, 478–489 (2006).

36. Dutta, R. et al. Demyelination causes synaptic alterations in hippocampi from multiple sclerosis patients. Ann. Neurol. 69, 445–454 (2011).

37. Drier, Y., Sheffer, M. & Domany, E. Pathway-based personalized analysis of cancer. Proc. Natl. Acad. Sci. U. S. A. 110, 6388–93 (2013).

