## Supplementary Table 1 for "Bile acid metabolism is altered in multiple sclerosis and supplementation ameliorates neuroinflammation"

**Supplementary Table 1. List of bile acid metabolites detected by untargeted metabolomics analyses in the discovery cohort**

| Metabolite |
|---|
| Cholic Acid (CA) |
| Chenodeoxycholic Acid (CDCA) |
| Deoxycholic Acid (DCA) |
| Ursodeoxycholic Acid (UDCA) |
| Hyocholic Acid (HCA) |
| Glycocholic Acid (GCA) |
| Glycocholic Acid Sulfate (GCAS) |
| Glycochenodeoxycholic Acid (GCDCA) |
| Glycochenodeoxycholic Acid Sulfate (GCDCAS) |
| Glycochenodeoxycholic Acid Glucuronide (GCDCAG) |
| Glycodeoxycholic Acid (GDCA) |
| Glycodeoxycholic Acid Sulfate (GDCAS) |
| Glycodeoxycholic Acid Glucuronide (GDCAG) |
| Glycoursodeoxycholic Acid (GUDCA) |
| Glycolithocholic Acid Sulfate (GLCAS) |
| Glycohyocholic Acid (GHCA) |
| Taurocholic Acid (TCA) |
| Taurocholic Acid Sulfate (TCAS) |
| Taurochenodeoxycholic Acid (TCDCA) |
| Taurodeoxycholic Acid (TDCA) |
| Taurolithocholic Acid Sulfate (TLCAS) |

**Supplementary Table 2. Demographic characteristics of discovery and validation cohorts**

| Characteristic | HC | RRMS | PMS | p-value |
|---|---|---|---|---|
| ***Discovery cohort*** | ***n=52*** | ***n=56*** | ***n=51*** | |
| Age, mean (SD) | 46 (13) | 39 (10) | 56 (8) | < 0.0001 |
| Sex, F:M | 35:17 | 41:15 | 39:12 | 0.57 |
| Race, n (%) | | | | 0.4 |
| Caucasian | 41(79) | 45 (80) | 45 (88) | |
| AA | 8 (15) | 7 (13) | 6 (12) | |
| Other | 3 (6) | 4 (7) | - | |
| Disease duration, median (IQR) | - | 8 (5) | 25 (9) | |
| Treatment, n (%) | - | | | |
| None | | 15 (27) | 24 (47) | |
| Injectable | | 38 (68) | 22 (43) | |
| Oral | | - | 3 (6) | |
| High-potency | | - | 2 (4) | |
| ***Validation cohort*** | ***n=75*** | ***n=50*** | ***n=125*** | |
| Age, mean (SD) | 49 (14) | 49 (13) | 56 (11) | 0.0004 |
| Sex, F:M | 46:29 | 31:19 | 75:50 | 0.96 |
| Race, n (%) | | | | 0.14 |
| Caucasian | 58 (77) | 39 (78) | 106 (85) | |
| AA | 9 (12) | 7 (14) | 17 (14) | |
| Other | 8 (11) | 4 (8) | 2 (1) | |
| Disease duration, median (IQR) | - | 13 (6) | 18.5 (11.5) | |
| Treatment, n (%) | - | | | |
| None | | 9 (18) | 56 (45) | |
| Injectable | | 32 (64) | 34 (27) | |
| Oral | | 2 (4) | 14 (11) | |
| High-potency | | 6 (12) | 15 (12) | |

**Supplementary Table 3. List of metabolites detected by targeted metabolomics analysis with range of detection in the targeted cohort**

| Analyte | Calibration Ranges (ng/mL) | |
|---|---|---|
| | LLOQ | ULOQ |
| Cholic Acid (CA) | 2.50 | 1250 |
| Chenodeoxycholic Acid (CDCA) | 5.00 | 2500 |
| Deoxycholic Acid (DCA) | 5.00 | 2500 |
| Lithocholic Acid (LCA) | 2.50 | 1250 |
| Ursodeoxycholic Acid (UDCA) | 5.00 | 2500 |
| Glycocholic Acid (GCA) | 2.50 | 1250 |
| Glycochenodeoxycholic Acid (GCDCA) | 5.00 | 2500 |
| Glycodeoxycholic Acid (GDCA) | 2.50 | 1250 |
| Glycoursodeoxycholic Acid (GUDCA) | 5.00 | 2500 |
| Glycolithocholic Acid (GLCA) | 2.50 | 1250 |
| Taurocholic Acid (TCA) | 2.50 | 1250 |
| Taurochenodeoxycholic Acid (TCDCA) | 5.00 | 2500 |
| Taurodeoxycholic Acid (TDCA) | 5.00 | 2500 |
| Taurolithocholic Acid (TLCA) | 2.50 | 1250 |
| Tauroursodeoxycholic Acid (TUDCA) | 2.50 | 1250 |

**Supplementary Table 4. Demographic characteristics of pediatric cohort**

| **Characteristic** | **HC (n=31)** | **Pediatric-onset MS (n=31)** | **P value** |
|---|---|---|---|
| Age, mean (SD) | 14.2 (2.3) | 14.2 (2.3) | n.s |
| Sex (1:2) | 14:17 | 14:17 | n.s |
| Race, n (%) | | | |
| Caucasian | 29 | 22 | |
| AA | - | - | |
| Other | 2 | 9 | |
| Disease duration, median (IQR) | - | 2.4 (2.5) | |
| DMT use, n (%) | - | 23 (74) | |

**Supplementary Table 5. List of bile acid metabolites detected by untargeted metabolomics analyses in the pediatric cohort**

| Metabolite |
|---|
| Cholic Acid (CA) |
| Chenodeoxycholic Acid (CDCA) |
| Deoxycholic Acid (DCA) |
| Ursodeoxycholic Acid (UDCA) |
| Hyocholic Acid (HCA) |
| Glycocholic Acid (GCA) |
| Glycocholic Acid Sulfate (GCAS) |
| Glycochenodeoxycholic Acid (GCDCA) |
| Glycochenodeoxycholic Acid Sulfate (GCDCAS) |
| Glycochenodeoxycholic Acid Glucuronide (GCDCAG) |
| Glycodeoxycholic Acid (GDCA) |
| Glycodeoxycholic Acid Sulfate (GDCAS) |
| Glycoursodeoxycholic Acid (GUDCA) |
| Glycolithocholic Acid Sulfate (GLCAS) |
| Glycohyocholic Acid (GHCA) |
| Glyco-alpha Muricholic Acid (GAMCA) |
| Glyco-beta Muricholic Acid (GBMCA) |
| Taurocholic Acid (TCA) |
| Taurocholic Acid Sulfate (TCAS) |
| Taurochenodeoxycholic Acid (TCDCA) |
| Taurodeoxycholic Acid (TDCA) |
| Taurolithocholic Acid Sulfate (TLCAS) |
| Isoursodeoxycholic Acid (IUDCA) |

**Supplementary Table 6. Characteristics of MS patients providing autopsy tissue**

| Case ID | MS type | Sex | Age (yrs) | Disease Duration (yrs) | Final EDSS | Post-mortem interval (hrs) |
|---|---|---|---|---|---|---|
| MS115 | SPMS | M | 67 | 25 | 8 | 11.25 |
| MS169 | SPMS | M | 35 | 21 | 9.5 | 9.5 |

### Supplementary Table 7. List of primers for murine astrocyte and microglial qPCR

| Name | Forward Primer | Reverse Primer |
|---|---|---|
| Amigo2 | GAGGCGACCATAATGTCGTT | GCATCCAACAGTCCGATTCT |
| Arg1 | TTTTAGGGTTACGGCCGGTG | CCTCGAGGCTGTCCTTTTGA |
| b actin | ACCTTCTACAATGAGCTGCG | CTGGATGGCTACGTACATGG |
| C1q | TCTGCACTGTACCCGGCTA | CCCTGGTAAATGTGACCCTTTT |
| C3 | AGCTTCAGGGTCCCAGCTAC | GCTGGAATCTTGATGGAGACGC |
| Cd14 | GGACTGATCTCAGCCCTCTG | GCTTCAGCCCAGTGAAAGAC |
| Clcf1 | CTTCAATCCTCCTCGACTGG | TACGTCGGAGTTCAGCTGTG |
| Cp | TGTGATGGGAATGGGCAATGA | AGTGTATAGAGGATGTTCCAGGTCA |
| Fkbp5 | TATGCTTATGGCTCGGCTGG | CAGCCTTCCAGGTGGACTTT |
| Gbp2 | GGGGTCACTGTCTGACCACT | GGGAAACCTGGGATGAGATT |
| Ggta1 | GTGAACAGCATGAGGGGTTT | GTTTTGTTGCCTCTGGGTGT |
| H2-D1 | TCCGAGATTGTAAAGCGTGAAGA | ACAGGGCAGTGCAGGGATAG |
| H2-T23 | GGACCGCGAATGACATAGC | GCACCTCAGGGTGACTTCAT |
| Iigp1 | GGGGCAATAGCTCATTGGTA | ACCTCGAAGACATCCCCTTT |
| Il1a | CGCTTGAGTCGGCAAAGAAAT | CTTCCCGTTGCTTGACGTTG |
| NOS2 | GCAAACATCACATTCAGATCCC | TCAGCCTCATGGTAAACACG |
| Lcn2 | CCAGTTCGCCATGGTATTTT | CACACTCACCACCCATTCAG |
| Psmb8 | CAGTCCTGAAGAGGCCTACG | CACTTTCACCCAACCGTCTT |
| Ptx3 | AACAAGCTCTGTTGCCCATT | TCCCAAATGGAACATTGGAT |
| Serping1 | ACAGCCCCCTCTGAATTCTT | GGATGCTCTCCAAGTTGCTC |
| S1pr3 | AAGCCTAGCGGGAGAGAAAC | TCAGGGAACAATTGGGAGAG |
| SIc10a6 | GCTTCGGTGGTATGATGCTT | CCACAGGCTTTTCTGGTGAT |
| Sgrn | GCAAGGTTATCCTGCTCGGA | TGGGAGGGCCGATGTTATTG |
| Steap4 | CCCGAATCGTGTCTTTCCTA | GGCCTGAGTAATGGTTGCAT |
| Timp1 | AGTGATTTCCCCGCCAACTC | GGGGCCATCATGGTATCTGC |
| Tm4sf1 | GCCCAAGCATATTGTGGAGT | AGGGTAGGATGTGGCACAAG |
| Tnfa | TGTGCTCAGAGCTTTCAACAA | CTTGATGGTGGTGCATGAGA |

### Supplementary Table 8. List of antibodies utilized for IHC of MS and EAE tissue

| Antibody | Manufacturer | Catalog # | Clone | Isotype | Host | Dilution |
|---|---|---|---|---|---|---|
| GFAP | Dako | GA52461-2 | Polyclonal | IgG | Rabbit | 1:1000 |
| GFAP | Cell Signaling Technology | 3670 | Mono | IgG1 | Mouse | 1:250 |
| Iba-1 | Wako | 019-19741 | Polyclonal | | Rabbit | 1:300 |
| iNos | Santa Cruz Biotechnology | sc-7271 | Mono; NOS2(C-11) | IgG1 | Mouse | 1:1000 |
| CD3 | Dako | A045201-2 | Polyclonal | | Rabbit | 1:200 |
| Mac-2 | BioLegend | 125401 | Mono; M3/38 | IgG2a | Rat | 1:200 |
| PSMB8 | Invitrogen | MA5-15890 | Mono; 1A5 | IgG1 | Mouse | 1:200 |
| GPBAR1 | Abcam | ab72608 | Polyclonal | IgG | Rabbit | 1:100 |
| FXR | Perseus Proteomics Inc | PP-A9033A-00 | Monoclonal | IgG | Mouse | 1:250 |
| GPBAR1 | Thermo Fisher Scientific | PA5-27076 | Polyclonal | IgG | Rabbit | 1:250 |
| GFAP | Dako | Z0334 | Polyclonal | IgG | Rabbit | 1:1000 |

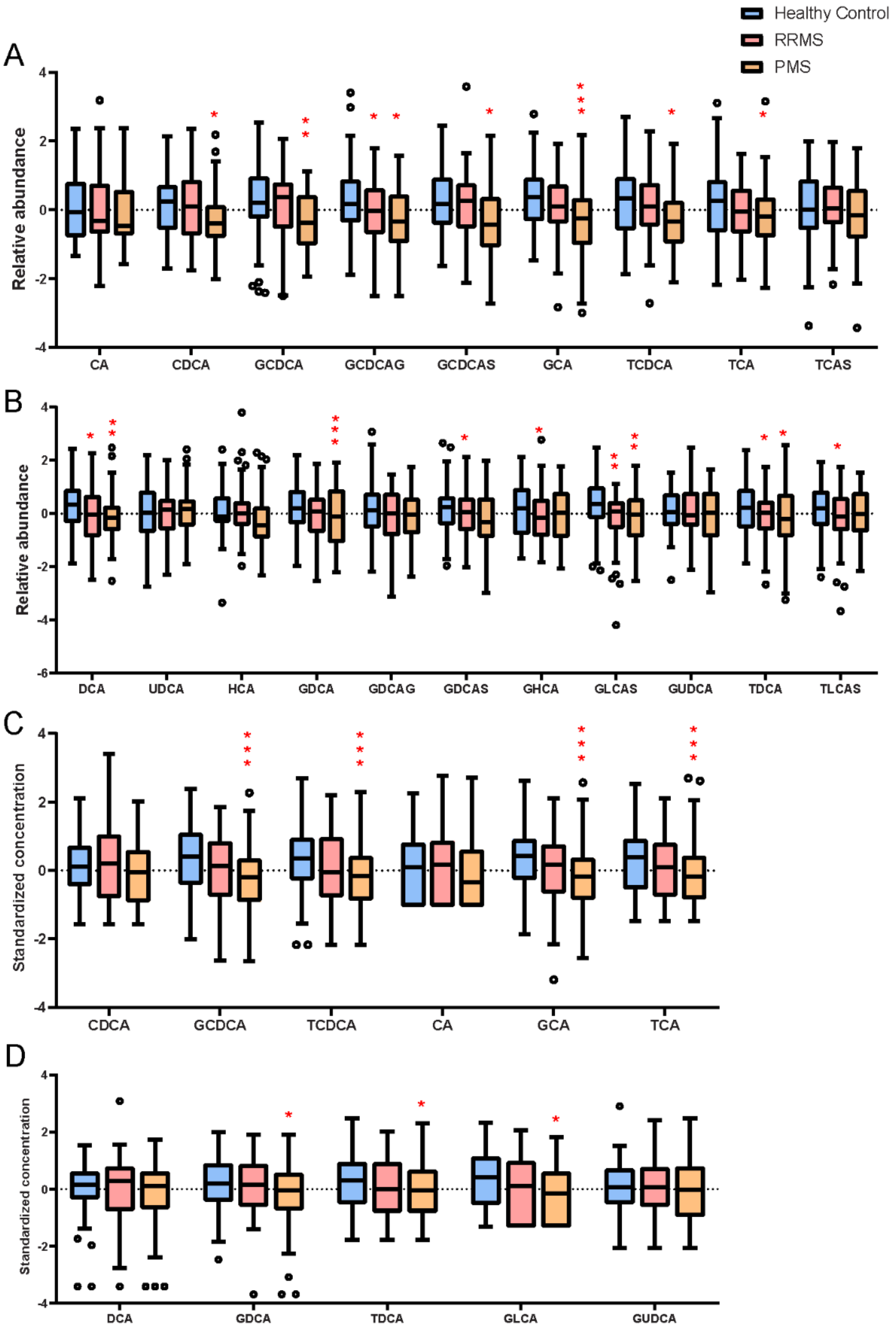


**Supplementary Figure 1. Individual bile acid metabolites from adult cohorts**

Box plots of primary bile acid metabolite relative abundances in the discovery cohort are shown in **(a)**, while relative abundances of secondary bile acid metabolites are shown in **(b)**. **(c)** Standardized concentrations of primary bile acids in the validation cohort, while **(d)** depicts standardized concentrations of secondary bile acids in the validation cohort. Red asterisks depict statistical significance of comparison to the control group with p values derived from multivariate regression models adjusting for age, sex and race (* $p<0.05$, ** $p<0.01$, *** $p<0.005$).

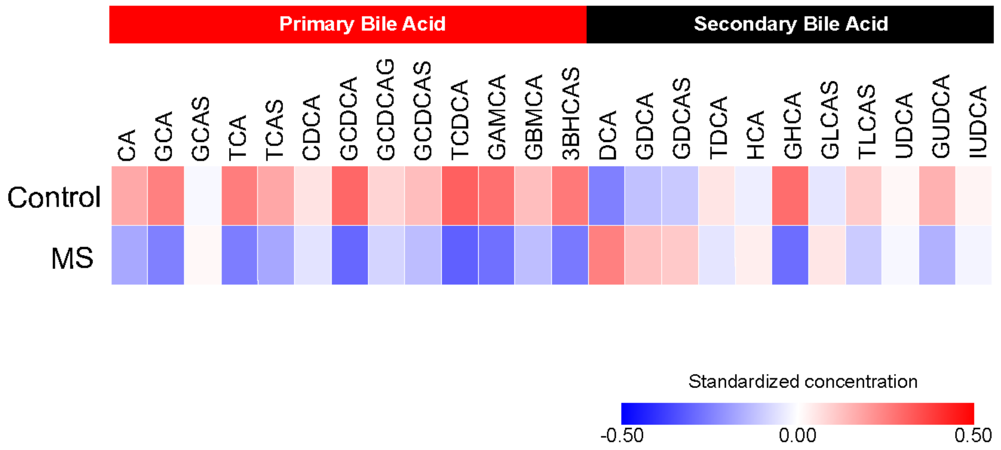


**Supplementary Figure 2. Heatmap of bile acid metabolite abundance from the pediatric cohort**

Heat map of mean standardized relative abundance of various bile acid metabolites (primary and secondary bile acid metabolites) identified in the circulation of pediatric-onset MS patients and healthy controls. CA – cholic acid, GCA – glycocholic acid, GCAS – glychocholic acid sulfate, TCA – taurocholic acid, TCAS – taurocholic acid sulfate, CDCA – chenodeoxycholic acid, GCDCA – glycochenodeoxycholic acid, GCDCAG – glycochenodeoxycholic acid glucuronide, GCDCAS – glycochenodeoxycholic acid sulfate, TCDCA – taurochenodeoxycholic acid,  GAMCA – glycoalpha-muricholic acid, GBMCA – glycobeta-muricholic acid, 3BHCAS – 3-beta hydroxy-5-cholenoic acid, DCA – deoxycholic acid, GDCA – glycodeoxycholic acid, GDCAS – glycodeoxycholic acid sulfate, TDCA -taurodeoxycholic acid, HCA -hyocholic aicd, GHCA - glycohyocholic acid, GLCAS -glycolithocholic acid sulfate, TLCAS – taurolithocholic acid sulfate, UDCA – ursodeoxycholic acid, GUDCA – glycoursodeoxycholic acid, IUDCA – isoursodeoxycholic acid.

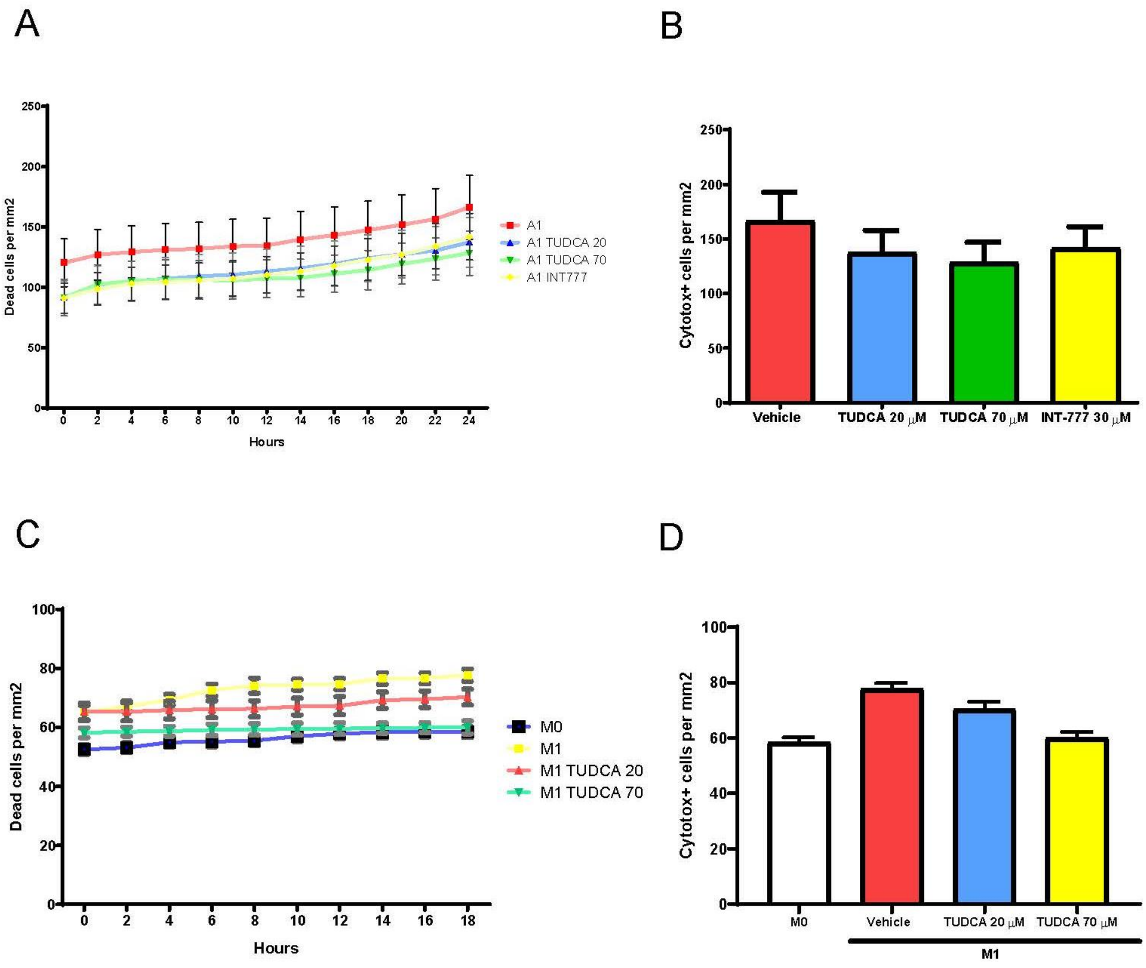


**Supplementary Figure 3. TUDCA treatment does not adversely affect the viability of murine astrocytes or microglia**

**(a)** We assessed viability of astrocytes, over a 24-hour period, in various culture conditions – A1 polarizing conditions (IL-1α, TNF-α and C1q) plus vehicle or varying doses of TUDCA or INT-777, using labelling with a cell viability dye and noted no increase in cell death with the addition of TUDCA or INT-777. **(b)** Quantification of cell death at the 24-hour time point. **(c)** We performed similar assessments, over an 18-hour period, in microglial cultures – either M0 condition or M1 polarization (IFN-γ and LPS) either with vehicle or varying doses of TUDCA and noted no increased cell death in the TUDCA conditions compared to vehicle. **(d)** Quantification of cell death at the 18-hour time point. Error bars in A-D represent standard error of the mean.

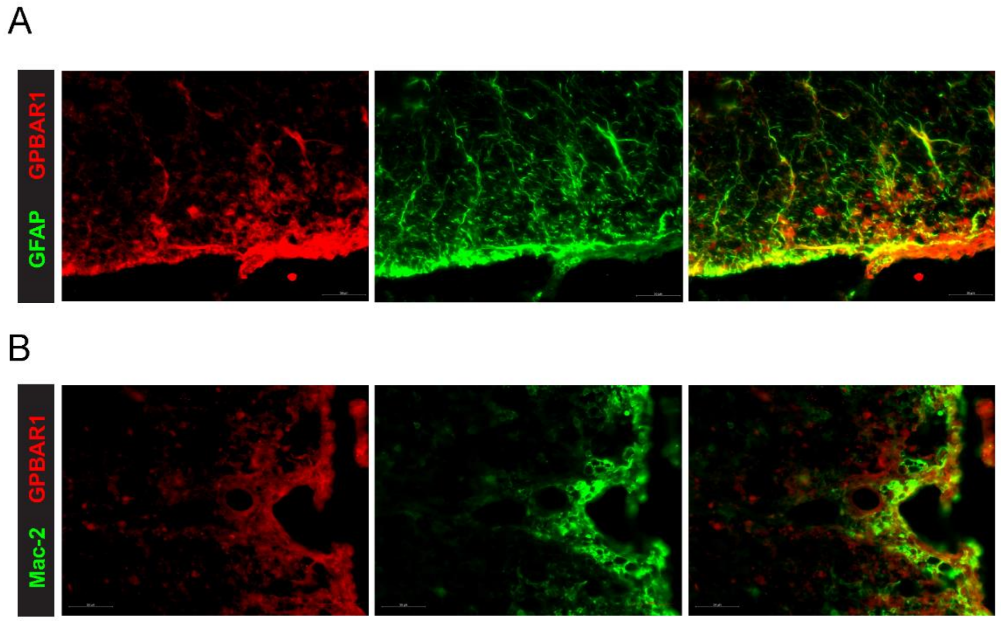


**Supplementary Figure 4. Bile acid receptors are found on astrocytes and microglia/macrophages in EAE**

**(a)** Immunohistochemistry for GFAP and GPBAR1 on spinal cord sections from mice with EAE, demonstrates the presence of several GPBAR1+ GFAP+ astrocytes. **(b)** We also performed immunohistochemistry for Mac-2 which stains myeloid cells and GPBAR-1, which revealed the presence of several GPBAR-1+ Mac-2+ cells in the spinal cord of EAE mice.

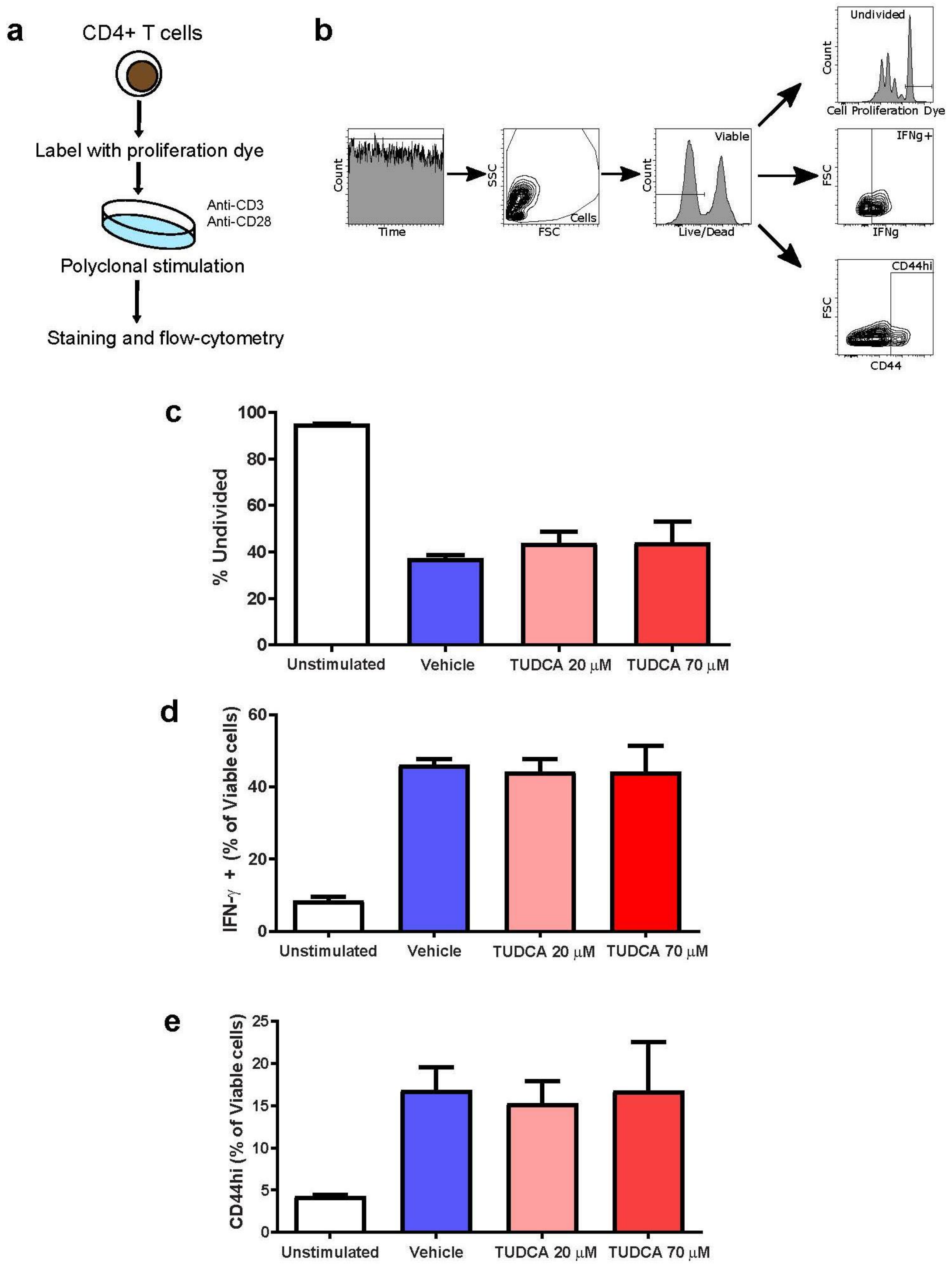


**Supplementary Figure 5. TUDCA treatment does not affect murine T cell proliferation and cytokine production**

**(a)** Murine CD4+ T cells isolated using negative bead selection from mouse splenocytes were stained with cell proliferation dye and then cultured in c-RPMI and polyclonally stimulated with anti-CD3 and anti-CD28 antibodies, in the presence or absence of varying doses of TUDCA. Following 72 hours of stimulation, we performed flow cytometry (gating strategy depicted in **(b)** and noted no significant effect of low- or high-dose TUDCA on T cell proliferation compared to vehicle **(c)**. We also noted no significant effects of TUDCA treatment on **(d)** interferon-gamma production or **(e)** T cell activation – based on CD44 expression ($CD44^{hi}$). Data is derived from a representative experiment. Error bars in b-d represent standard error of the mean.
